# The Gut Microbiome of Exudivorous Marmosets in the Wild and Captivity

**DOI:** 10.1101/708255

**Authors:** Joanna Malukiewicz, Reed A. Cartwright, Jorge A. Dergam, Claudia S. Igayara, Sharon Kessler, Silvia B. Moreira, Leanne T. Nash, Patricia A. Nicola, Luiz C.M. Pereira, Alcides Pissinatti, Carlos R. Ruiz-Miranda, Andrew T. Ozga, Adriana A. Quirino, Christian Roos, Daniel L. Silva, Anne C. Stone, Adriana D. Grativol

## Abstract

Mammalian captive dietary specialists like folivores are prone to gastrointestinal distress and primate dietary specialists suffer the greatest gut microbiome diversity losses in captivity compared to the wild. Marmosets represent another group of dietary specialists, exudivores that eat plant exudates, but whose microbiome remains relatively less studied. The common occurrence of gastrointestinal distress in captive marmosets prompted us to study the *Callithrix* gut microbiome composition and predictive function through bacterial 16S ribosomal RNA V4 region sequencing. We sampled 59 wild and non-wild *Callithrix* across four species and their hybrids. Host environment had a stronger effect on the gut microbiome than host taxon. Wild *Callithrix* gut microbiomes were enriched for *Bifidobacterium*, which process host-indigestible carbohydrates. Captive marmoset guts were enriched for Enterobacteriaceae, a family containing pathogenic bacteria. While gut microbiome function was similar across marmosets, Enterobacteriaceae seem to carry out most functional activities in captive host guts. More diverse bacterial taxa seem to perform gut functions in wild marmosets, with *Bifidobacterium* being important for carbohydrate metabolism. Non-wild marmosets showed gut microbiome composition aspects seen in human gastrointestinal diseases. Thus, captivity may perturb the exudivore gut microbiome, which raises implications for captive exudivore welfare and calls for husbandry modifications.

## Introduction

The mammalian gut microbiome plays an important role in host physiology^1,2^, and microbiome dysbiosis is thought to negatively impact host health^3–5^. More closely related hosts seem to share more similar microbiome communities than more distantly related hosts (i.e., phylosymbiosis)^6,7^, and gut microbiome communities are usually enriched for bacteria associated with the main macronutrients of a host’s feeding strategy^8–12^. Yet, environmental factors significantly alter individual host microbiomes^10,12^, as evidenced by differences in microbiome composition between wild and captive conspecifics across a variety of animal taxa^13–19^. Gut microbiome studies of captive and wild mammals show that non-human primates (NHPs) experience relatively large losses of native gut microbiome diversity in captivity compared to the wild^5,13^. Additionally, dietary specialist NHPs including folivores (leaf-eating) and frugo-folivores (fruit and leaf-eating) are especially prone to gastrointestinal problems in captivity^20–24^. Among humans and NHPs, dysbiosis in gut microbiome composition has been tied to gastrointestinal diseases^4,22,25^.

A number of mammals, including some primates, are exudivorous, meaning that they nutritionally exploit viscous plant exudates that are composed of polysaccharides^26,27^ such as galactan, mannose, arabianans, arabinose, xylose, and glucuronic acid (e.g.,^28–30^). Among mammalian dietary specialists, the exudivore gut microbiome remains relatively little studied. Nonetheless, Brazilian *Callithrix* marmosets, a relatively recent genus of closely-related NHP exudivores^31^, are excellent models to study exudivore gut microbiomes. In the wild, these primates nutritionally exploit hard to digest oligosaccharides of tree gums or hardened saps that require fermentation by gut microbioata for digestion^32,33^. Host specific gastrointestinal adaptions in marmosets that likely facilitate microbial polysaccharide fermentation include an enlarged cecum, an elongated colon, and gut transit times attuned to gum digestion^34–36^. Further, *Callithrix* species collectively possess a number of morphological adaptations in cranial shape and musculature, dentition, and nail shape that allow them to access natural gum sources by gouging and scraping hard plant surfaces such as bark^37–39^.

Marmosets are regularly maintained in captivity as biomedical research models, for captive breeding of endangered *C. aurita*, and due to illegal pet trafficking^31^. In captivity, marmosets commonly develop symptoms of gastrointestinal distress like inflammatory-like bowel disease, chronic malabsorption, chronic diarrhea, chronic enteritis, and chronic colitis without clear pathogenesis^40–42^. Up to now, most *Callithrix* gut microbiome studies have focused on captive *C. jacchus* to identify specific bacterial strains and on how life history, social, or laboratory conditions affect the gut microbiome composition^41^. A review of these studies suggests that there may be an association between gastrointestinal distress and gut microbiome dysbiosis in *Callithrix*^41^. A necessary first step towards understanding diseased gut microbiome composition profiles is defining baseline gut microbiome composition variation and function of non-diseased individuals^2^. Thus, comparing the gut microbiome of wild and captive conspecifics is an important step for such approaches.

Here, we determine gut microbiome profiles of *Callithrix* sampled in and out of captivity throughout Brazil. We applied 16S ribosomal RNA (rRNA) V4 region amplicon sequencing of *Callithrix* gut microbiota, and investigated gut microbiome composition and gut microbiome predictive functional profiles. Anal swabs were sampled in Brazil from 59 healthy individuals of four species and three hybrid types (Table 1) that were either wild, translocated into captivity from the wild, or born into captivity (Figure 1). Our specific aims in this study were to evaluate the influence of host taxon and environment on *Callithrix* gut microbiome composition, diversity, and function. As marmosets are considered obligate exudivores^27^, we hypothesize that marmoset gut microbiome composition and predictive functional profiles are strongly biased toward carbohydrate metabolism across marmoset taxa. Yet, as previous studies have shown differences in gut microbiome composition between wild and captive animal hosts^5,13^, we hypothesize that *Callithrix* gut microbiome composition between individual hosts differs according to host environmental status (ie., captive, translocated, wild).

**Table 1.** Information summary on marmoset host taxon, sampling location, hybrid status, sampling location and environment. For sampling locations, the following abbreviations are used: CPRJ= Centro de Primatologia do Rio de Janeiro, CEMAFAUNA= Centro de Conservação e Manejo de Fauna da Caatinga, and Setor de Etologia, SERCAS=Reintrodução e Conservação de Animais Silvestres. For host environment, the following abbreviations are used: W=Wild, T= Translocated, C= Captive.

| Host Taxon | Sampling Location | Approximate Collection Geographic Coordinates | N | Host Environment |
| --- | --- | --- | --- | --- |
| *C. aurita* | Guiricema, Minas Gerais, Brazil | -21.008, -42.723 | 2 | W |
| *C. aurita* | CPRJ, Guapimirim, Rio de Janeiro, Brazil<br>(wild marmosets originally from Natividade, Rio de Janeiro, Brazil) | -21.061, -41.977 | 3 | T |
| *C. aurita* | CPRJ, Guapimirim, Rio de Janeiro, Brazil | -22.4881, -42.913 | 5 | C |
| *C. aurita* x *Callithrix* sp. | CPRJ, Guapimirim, Rio de Janeiro, Brazil | -22.489, -42.914 | 1 | C |
| *C. geoffroyi* | CPRJ, Guapimirim, Rio de Janeiro, Brazil | -16.931, -42.4852 | 3 | C |
| *C. geoffroyi* | Berilo, Minas Gerais, Brazil | -22.489, -42.914 | 1 | W |
| *C. jacchus* | Guarulhos Municipal Zoo, Guarulhos, São Paulo, Brazil | -23.443, -46.554 | 9 | C |
| *C. penicillata* | Guarulhos Municipal Zoo, Guarulhos, São Paulo, Brazil | -23.443, -46.554 | 4 | C |
| *C. penicillata* | CPRJ, Guapimirim, Rio de Janeiro, Brazil | -22.486, -42.914 | 1 | T |
| *C. penicillata* | CEMAFAUNA, Petrolina, Pernambuco, Brazil | -9.327, -40.544 | 2 | C |
| *C. jacchus* x *C. penicillata* | Guarulhos Municipal Zoo, Guarulhos, São Paulo, Brazil | -23.443, -46.554 | 1 | C |
| *C. jacchus* x *C. penicillata* | SERCAS, Campos, RJ, Brazil<br>(wild marmosets originally from Ilha D'Água, Rio de Janeiro, RJ, Brazil) | -22.810, -43.163 | 16 | T |
| *C. jacchus* x *C. penicillata* | CPRJ, Guapimirim, Rio de Janeiro, Brazil | -22.490, -42.914 | 6 | T |
| *C. penicillata* x *C. geoffroyi* | Viçosa, Minas Gerais, Brazil | -20.764, -42.900 | 5 | W |

**Figure 1.**
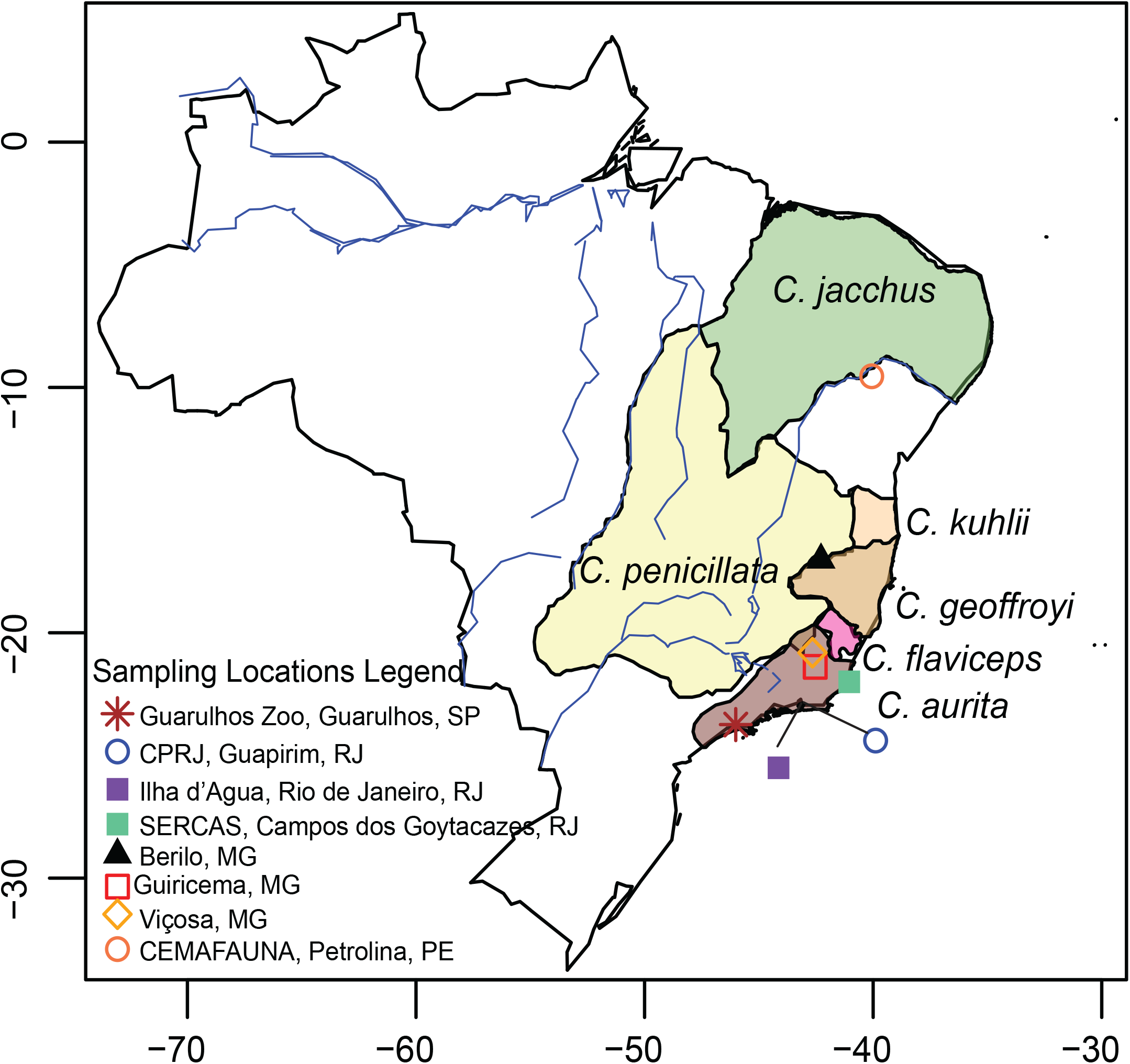
Natural *Callithrix* ranges and sampling locations in Brazil. Sampling locations are represented by different colored shapes and species names are written next to their respective ranges. Ranges are based on 2012 IUCN Red List Spatial Data from http://www.iucnredlist.org/technical-documents/spatial-data. Legend abbreviations are as follows-MG: Minas Gerais, Rio de Janeiro: RJ, Pernambuco: PE, São Paulo: SP; CPRJ: Centro de Primatologia do Rio de Janeiro; CEMAFAUNA: Centro de Conservação e Manejo da Fauna da Caatinga.

## Methods

### Sample Collection

We collected anal swabs between 2015 and 2016 from 59 adult individuals, and general sampling information is summarized in Table 1 and Figure 1. Supplementary Table S1 gives detailed information on each sampled marmoset host including taxon, sampling location, and environment. We considered marmosets older than 11 months as adults, following age criteria based on dental characteristics and genitalia growth^43^. Marmoset sampling was authorized and approved by the Brazilian Environmental Ministry (SISBIO protocol# 47964-2), and the Arizona State University IACUC (protocol# 15-144R). Wild animals were captured with Tomahawk style traps baited with bananas. As part of a larger marmoset ‘omics’ study (e.g.^31^), sampled animals were immobilized with ketamine (10 mg/kg of body weight) through inner thigh intramuscular injection, photographed, weighed, measured, examined clinically by veterinarians, and deemed healthy upon examination. Copan FLOQ Swabs were gently rotated in the anal region and submerged in storage buffer (50 mM Tris pH 8.0, 50 mM EDTA, 50 mM Sucrose, 100 mM NaCl, 1% SDS) before being discarded. After processing, animals were returned to cages for recovery. Wild marmosets were released at original capture sites. Host taxon identification followed previously published phenotype descriptions^44,45^ and personal observations by JM and CSI. Hosts were also classified by their environment as wild (captured as free-range individuals), translocated (born wild but later put into captivity), or captive (born and raised in captivity). This study is reported in accordance to ARRIVE guidelines (https://arriveguidelines.org/resources/questionnaire). All methods were carried out in accordance with relevant international guidelines and regulations.

### Sample Processing and Sequencing

Bacterial DNA extraction from *Callithrix* anal swabs was carried out by following a modified phenol-chloroform protocol^46^. Modifications included beating the samples on a vortex fitted with a horizontal vortex adaptor (#13000-V1-24, Mo Bio, Carlsbad, CA, USA) for 10 minutes at step “2Aiii,” precipitating samples in 100% ethanol in step “2Axvi” and rehydrating DNA pellets in 25 uL low TE buffer at step “2Axxii.” Extracted DNA was quantified on a Qubit3 (Life Technologies, Carlsbad, CA, USA) with a dsDNA HS Assay Kit (Life Technologies). DNA samples obtained for this study have been registered in the Brazilian SISGen database under entries # A2E885E, A965629, A5CB6FA, AE784B5, and A07A291. The V4 region of the bacterial 16S rRNA gene was amplified from sampled DNA in triplicate using the barcoded primer set 515f/806r^47^. Amplicon triplicates were combined for each individual and then pooled in equimolar amounts into a multiplexed Illumina sequencing library. The library was purified with a Zymo DNA Concentrator and Cleaner-5 (#D4013, Zymo Research, Irving, CA, USA) and size selected for 375-380 base pairs with Agencourt Ampure XP (#A63880, Beckman Coulter, Indianapolis, IN, USA) magnetic beads. Libraries were sequenced at Arizona State University, USA on an Illumina MiSeq for2×250 cycles.

### Bioinformatics and Statistical Analysis

Code for bioinformatics analysis described below is available at github.com/Callithrix-omics/callithrix_microbiome. Data were demultiplexed using default parameters in QIIME2-2021.2^48^. The DADA2 QIIME2 plug-in^49^ was used to quality-filter and trim sequences and join paired-end reads. Upon trimming, the first 10 and last 30 nucleotides were removed from reverse reads due to low base quality. These steps resulted in feature tables of DNA sequences and their per-sample counts. MAAFT^50^ and FastTree^51^, as part of the QIIME2 phylogeny plug-in, aligned and produced a mid-pointed rooted phylogenetic tree of feature sequences. Taxonomic composition of samples was determined with the QIIME2 Naive Bayes q2-feature-classifier plug-in, which was trained on pre-formatted SILVA reference sequence and taxonomy files “Silva 138 SSURef NR99 515F/806R region sequences” and “Silva 138 SSURef NR99 515F/806R region taxonomy”^52–54^for the portion of the 16S V4 region bounded by the 515F/806R primer pair. The pre-formatted files were downloaded from docs.qiime2.org/2021.4/data-resources. Taxonomic classification of the feature table was carried out with the q2-feature-classifier classify-sklearn command. For further down stream analyses, we used the QIIME2 export option to extract a biom format file from the classified feature table as well as feature table taxonomic information. Information from the exported biom file and feature table taxonomy were merged into a new biom format file with the biom 2.1.10^55^ command line tool.

For community profiling and comparative analysis, we used the ‘Marker-gene Data Profiling’ (MDP) module of the MicrobiomeAnalyst web-based platform^56^, using the merged biom file from above as well as sample metadata given in Supplementary Table S1. At the MicrobiomeAnalyst data filtering step, we left the default settings of the ‘Low count filter’ to a minimum count of 4 and 20% prevalence in samples and the ‘percentage to remove’ option under ‘Low variance filter’ set to 10% based on the interquantile range. Next, at the data normalization step, we chose to rarefy the data to the minimum library size, data was scaled by ‘total sum scaling,’ and we did not apply any data transformations. Marmoset gut microbiome richness (ie., the number of observed host gut microbiome features as determined in QIIME) was calculated as a measure of alpha-diversity in the MicrobiomeAnalyst ‘Alpha-Diversity Analysis’ submodule. Normalized and filtered data were used in the module with settings configured for host taxon and environment, respectively, and ‘feature’ as the taxonomic level. We evaluated the relationship of marmoset gut microbiome alpha diversity with both host environment and taxon by fitting a Poisson distributed generalized linear model (GLM) in R^57^. In this GLM, host gut microbiome richness was set as the response variable, and host taxon and environment were used as the two independent variables. No interaction term was included in the GLM, as we assumed the effects of host taxon and environment were independent of each other. Analysis of deviance was used to determine the statistical significance of the inclusion of both independent variables in the fitted GLM. Model validity and fit was assessed with a plot of standardized deviance residuals against fitted values, Q-Q plot of quantile residuals, and identification of influential observation based on leverage and Cook’s distance. Post-hoc analyses for this model were performed with Tukey’s HSD test using the glht function from the multcomp^58^ R package.

To explore beta diversity of the *Callithrix* gut microbiome, we calculated the Bray-Curtis dissimilarity indices for each host, and then used the indices to make a non-metric multidimensional scaling (NMDS) ordination plot in the R vegan program^59^. We superimposed both environmental and taxon information for each marmoset on to the NMDS plot. To understand whether host environment and taxon had an effect on marmoset gut microbiome Bray-Curtis dissimilarity indices, we used adonis2 function in the phyloseq package^60^. We fitted PERMANOVA^61^ models which included the marginal effects of host environment and taxon as independent variables and Bray-Curtis dissimilarity indices as the dependent variable. Simulation studies have found that PERMANOVA is robust to unbalanced sampling designs^62^. The PERMANOVA model was ran with the adnois2 function. PERMANOVA post-hoc tests of Bray-Curtis dissimilarity indices were carried out as pairwise adonis tests with the adonis.pair function from the the EcolUtils^63^ R package. The test was run for 1000 permutations and p-values were corrected by the false discovery rate (FDR).

To profile gut microbiome bacterial taxa abundance, we used the’ Stacked Bar/Area Plot’ submodule of MicrobiomeAnalyst to generate stacked bars of relative bacterial abundance at various taxonomic levels (class and genus) according to host taxon and captivity, respectively. Taxa resolution settings were set to merge small taxa with total counts of less than 10. Average percentages of gut bacterial classes for marmosets according to host taxon and environment were calculated with the MicrobiomeAnalysis ‘Interactive Pie Chart Exploration’ submodule with same setting as for relative bacterial abundance. To test for significance in differential bacterial taxa abundance according to host environment and taxon, respectively, we used LEfSe^64^ at the class and genus level for bacterial taxa. The LEfSe submodule within MicrobiomeAnalyst was used with the default settings of a FDR-adjusted p-value cutoff set to 0.1 and the log LDA cut-off at 2.0.

To explore the functional aspects of the *Callithrix* gut microbiome, the Kyoto Encyclopedia of Genes and Genome Orthology (KEGG) pathways were predicted with PICRUSt2^65^ by following guidelines at https://github.com/picrust/picrust2/wiki. First, predicted KEGG ORTHOLOGY (KO) functional predictions were carried out with the metagenome_pipeline.py script with the –strat_out option. By default, PICRUSt2 excluded all features with the nearest sequenced taxon index (NSTI) value >2 from the output. The average weighted NSTI value of the data set after this automatic filtering was 0.08 ± 0.12 SD. Then Kyoto Encyclopedia of Genes and Genomes (KEGG) pathway abundances were derived from predicted KO abundances were performed with the “–no_regroup” option in the pathway_pipeline.py script in PICRUSt2. We then rounded the unstratified KEGG pathway abundance results for alpha and beta analysis of predicted functional pathways of the *Callithrix* gut microbiome. We then turned these results into a phyloseq object in R. For alpha diversity, we used phyloseq to estimate the observed number of *Callithrix* gut microbiome predicted KEGG pathways (i.e. the marmoset gut microbiome KEGG pathway richness). We then fit a GLM model with KEGG pathway richness in a similar manner as described above for marmoset gut microbiome composition analysis.

Using PICRUSt2 unstratified KEGG pathway abundance results, we generated a relative abundance plot of *Callithrix* gut microbiome KEGG metabolic processes using the Shotgun Data Profiling Module in MicrobiomeAnalyst. At the Microbiome-Analyst data filtering step, we left the default settings of the ‘Low count filter’ to a minimum count of 4 and 20% prevalence in samples and the ‘percentage to remove’ option under ‘Low variance filter’ set to 10% based on the interquantile range. After MicrobiomAnalyst filters, a total of 137 KEGG pathways remained for further analysis. A functional diversity relative abundance plot was generated for KEGG metabolism based on category abundance total hits. We grouped this abundance plot by first by host environment and then indicated host taxon for each host. We tested for significant patterns of differential abundance between host environment and taxon, respectively, in MicrobiomeAnalyst using the LEfSe submodule with a FDR-adjusted p-value cutoff of 0.1 and Log LDA score of 2.0. Functions of KEGG pathways were derived from the KEGG database^66^.

BURRITO^67^, an online interactive visualization module, was used to make links between our bacterial abundance data and predicted functional profiles from the *Callithrix* gut microbiome. As input, we used bacterial taxonomic abundance and taxonomy data based on the biom file originally extracted from QIIME2. We also provided a function attribution table based on PICRUSt2 output that linked the functional and taxonomic data by following instructions for the convert_table.py script at https://github.com/picrust/picrust2/wiki. We also provided a metadata table to BURRITO which included host environmental classifications. Host taxon information was later superimposed manually on result plots manually in Adobe Illustrator.

## Results

After initial processing and filtering of individual marmoset gut microbiome libraries, a total of 10,902,292 sequence reads was obtained with an average of 201,894 (124389.64 +/- SD) reads per sample. After quality filtering, 8,885,656 reads remained with an average 164,549.19 (99,524.230 +/- SD) reads per sample. Afterward, merging of paired-end sequences produced 8,191,034 reads, with an average of 151,685.81 (91,568.49 +/- SD) reads per sample. This information is detailed in Supplementary Table S2.

### Diversity of *Callithrix* Gut Microbiome Bacterial Taxa

Results of data rarefaction for *Callithrix* gut microbiome alpha diversity analyses are shown in Supplementary Figure S1. Boxplots of alpha diversity in terms of marmoset gut microbiome richness for host environment and taxonomic classification, respectively, are shown in Figure 2a-b. Individual host alpha diversity measures are listed in Supplementary Table S1. The GLM model fitted for the influence of host taxon and environment on marmoset gut microbiome alpha diversity is summarized in Table 2. In the model, post-hoc pairwise host environment comparisons between wild and translocated hosts as well as captive and translocated hosts were highly significant (Supplementary Table S3). For host taxon, respective post-hoc pairwise comparisons between *C. aurita* and *C. jacchus, C. penicillata*, and *C. jacchus* x *C. penicillata* were highly significant (Supplementary Table S3). Respective pairwise comparisons between *C. jacchus* and *C. geoffroyi* and *C. penicillata* x *C. geofforyi* were also significant (Supplementary Table S3).

**Figure 2.**
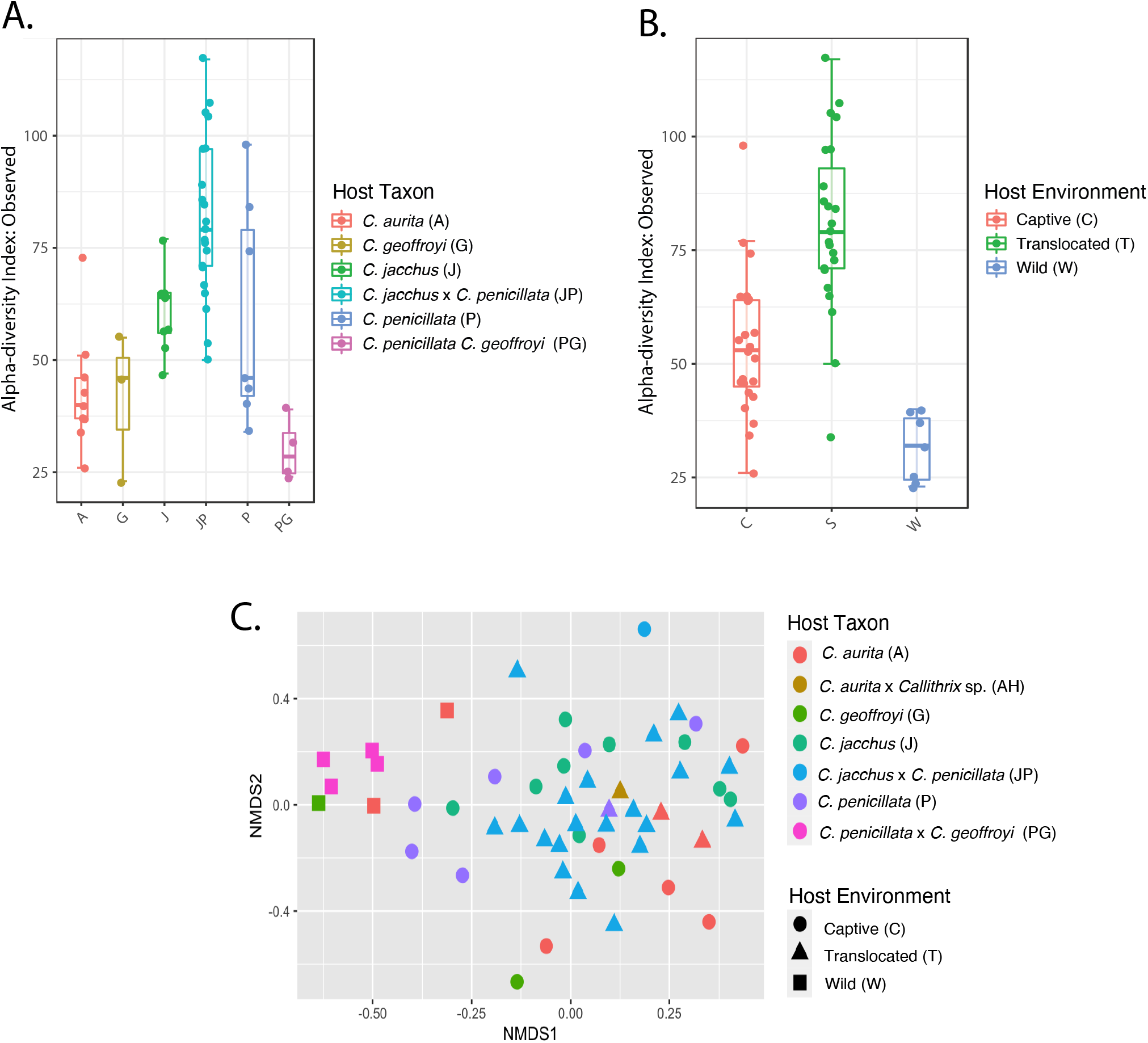
Boxplots of gut microbiome richness by host taxon (A) and index by host environment (B). Panel (C) shows an NMDS ordination plot for gut microbiome beta-diversity measured by the Bray-Curtis dissimilarity index.

**Table 2.** Analysis of deviance for GLM (Richness ∼ Host Taxon + Host Environment) fitted for *Callithrix* gut microbiome compositional alpha diversity.

| Term | Degrees of Freedom | Deviance | Residual Degrees of Freedom | Residual Deviance | p-value $\chi^2$ |
| --- | --- | --- | --- | --- | --- |
| Null |  |  | 52 | 483.32 |  |
| Host Taxon | 5 | 282.32 | 47 | 201.00 | < 2.20e-16 |
| Host Environment | 2 | 31.30 | 45 | 169.70 | 1.60e-07 |

For marmoset gut microbiome beta diversity, a NMDS plot of Bray-Curtis dissimilarity index with superimposed host environment and taxon is shown in Figure 2c. The effects of host environment on marmoset gut microbiome beta diversity were significant (PERMANOVA, R2=0.09, df=2, p=0.001), while those of host taxon were not (PERMANOVA, R2=0.14, df=6, p=0.06). Post-hoc analysis of all possible combinations of host environmental levels were found to be significant (p-value=0.001).

### *Callithrix* Gut Microbiome Bacterial Taxon Composition and Abundance

Figure 3a shows relative abundances of bacterial classes for hosts according to their environmental and taxon classification. These plots show that captive marmosets had relatively high abundance of Gammoproteobacteria (average abundance 60%). Translocated marmosets seem to have relatively high abundance of Campylobacteria (average abundance 36%). Wild marmosets have an average relative abundance of Campylobacteria of 33% and an Actinobacteria average relative abundance of 41%. For highest gut bacterial abundances among marmoset taxa, Gammaoproteobacteria was the most abundant bacterial class for *C. aurita* (39%), *C. geoffroyi* (55%), *C. jacchus* (72%), and *C. penicillata* (41%). Campylobacteria was most abundant in *C. jacchus* x *C. penicillata* hybrids (31%), while Actinobacteria was highest in *C. penicillata* x *C. geofforyi* hybrids (44%). Enterobacteriaceae were the most abundant bacterial family in the gut microbiome of captive marmosets (47%). For translocated marmosets, *Heliobacter* was most abundant in the gut microbiome (28%). Then for wild marmosets, the most abundant bacterial genus in the gut microbiome was *Bifidobacterium*. LefSe differential gut microbiome bacteria abundance analysis at class and genus levels support the statistical significance of these differences among marmoset hosts (Figure 3b-c, Supplementary Figure S2).

**Figure 3.**
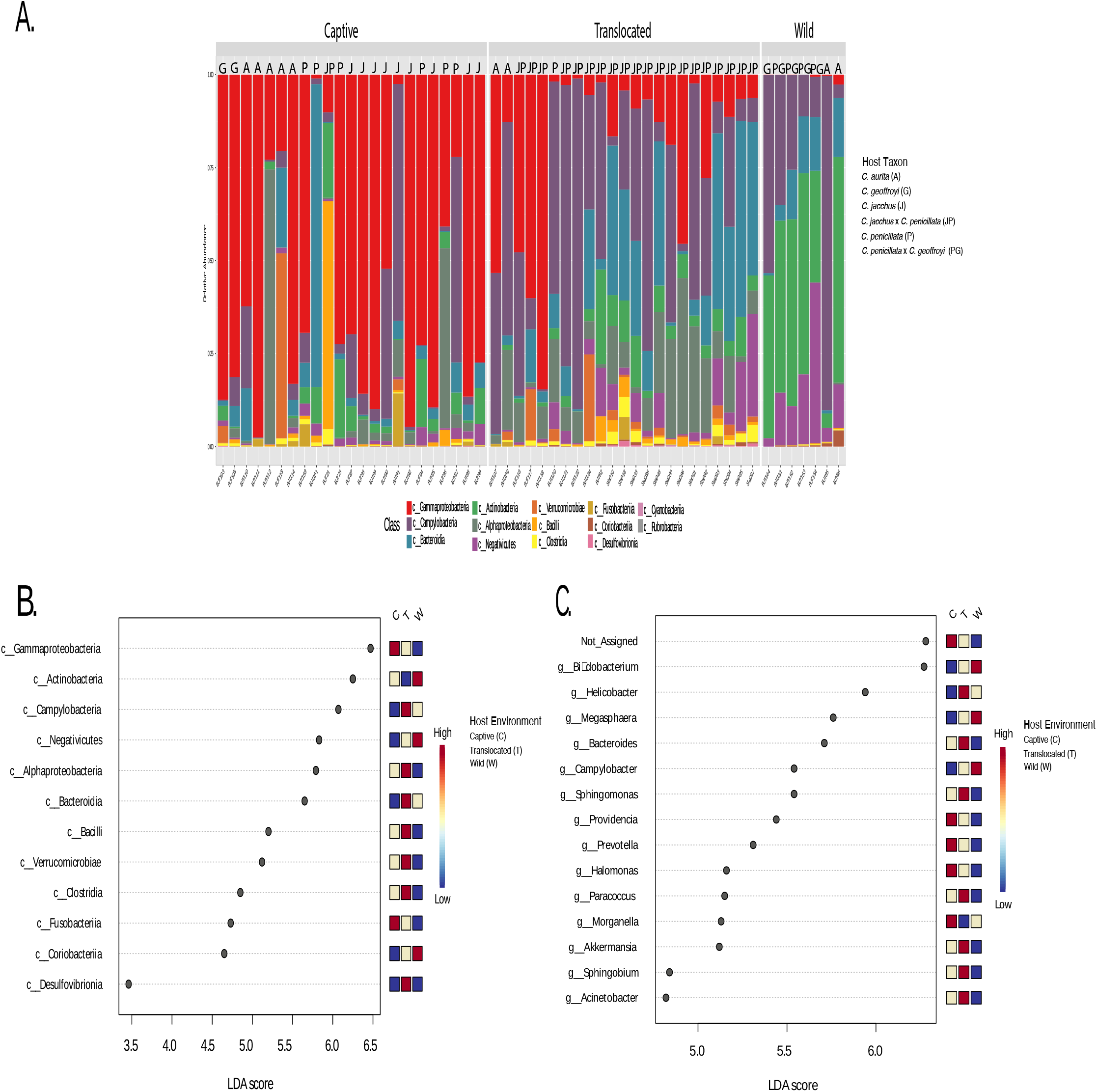
(A). Relative class level bacterial abundance (lower legend) by host environment (Captive, Translocated, and Wild) and taxon (see right-side legend). (B). LefSe analysis of bacterial class abundance categorized by host environment. (C). LefSe analysis of bacterial genus abundance categorized by host environment. The corresponding legends for plots B and C are to the right of both plots.

### Diversity of Predicted Functional Pathways of the *Callithrix* Gut Microbiome

A total of 183 KEGG predictive pathways were identified among our sampled marmoset hosts (Supplementary Table S4). Boxplots of KEGG pathways richness of the *Callithrix* gut microbiome according to host taxon and environment, respectively, are shown in Figure 4a-b. KEGG pathway richness values for individual hosts are listed in Supplementary Table S1. The GLM fit to explain the effects of host environment and taxon on gut KEGG pathway alpha diversity is summarized in Table 3. Neither host environment nor taxon were significant in the fitted model for having an effect on the alpha diversity of marmoset predicted KEGG pathways of the gut microbiome.

**Figure 4.**
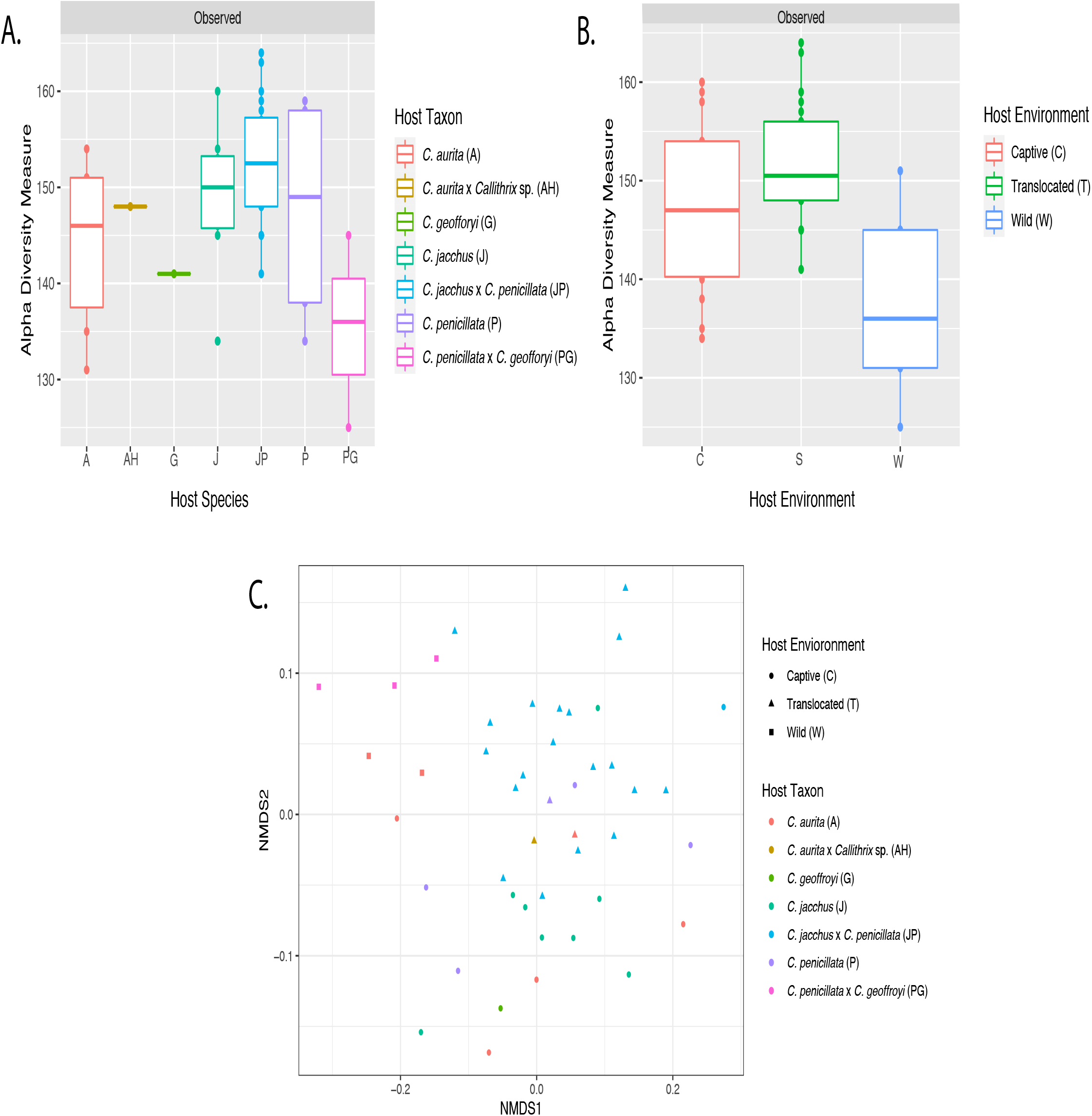
Boxplots of gut microbiome predicted gut KEGG pathways richness by host taxon (A) and host environment (B). Panel (C) shows an NMDS ordination plot for gut microbiome predicted gut KEGG pathways beta-diversity measured by the Bray-Curtis dissimilarity index. Legend of host classifications are shown on the right side of each plot.

**Table 3.** Analysis of deviance for GLM (Richness ∼ Host Taxon + Host Environment) fitted for *Callithrix* gut microbiome functional alpha diversity.

| Term | Degrees of Freedom | Deviance | Residual Degrees of Freedom | Residual Deviance | p-value $\chi^2$ |
| --- | --- | --- | --- | --- | --- |
| Null |  |  | 44 | 23.69 |  |
| Host Taxon | 6 | 7.30 | 38 | 16.39 | 0.29 |
| Host Environment | 2 | 0.18 | 36 | 16.21 | 0.92 |

For marmoset gut microbiome KEGG pathway beta diversity, Bray-Curtis dissimilarity index values were plotted on a NMDS ordination plot with superimposition of both host environment and taxon (Figure 4c). Neither the effects of host environment (PERMANOVA, R2=0.07, df=2 p=0.140) nor host taxon (PERMANOVA, R2=0.12, df=6, p=0.493) had a significant effect on *Callithrix* gut predicted KEGG pathway beta diversity.

### *Callithrix* Gut Microbiome KEGG Pathway Composition and Abundance

For relative abundance of predicted KEGG pathways of the *Callithrix* gut microbiome, sampled marmoset distributions showed an even distribution of KEGG metabolism categories, with carbohydrate metabolism being one of the most abundant categories (Figure 5a). For relative abundance of predicted KEGG pathways of the Callithrix gut microbiome, sampled marmoset distributions showed an even distribution of KEGG metabolism categories, with carbohydrate metabolism being one of the most abundant categories (Figure 5a). Visual inspection of the plot shows that this pattern holds regardless of host environmental or taxon categorization. LEfSe analysis for the top significantly enriched predicted KEGG pathways in the marmoset gut is shown for host environment in Figure 5b and for host taxon in Figure 5c. All top predicted KEGG gut microbiome pathways were enriched for in captive marmosets. The top most pathway was K01051 (LDA = 4.8) and is involved with carbohydrate metabolism of pectinesterase. This same pathway is also enriched in C. aurita and C. jacchus. The orthology of remaining pathways in Figure 5b-c is given in Supplementary Table S5.

**Figure 5.**
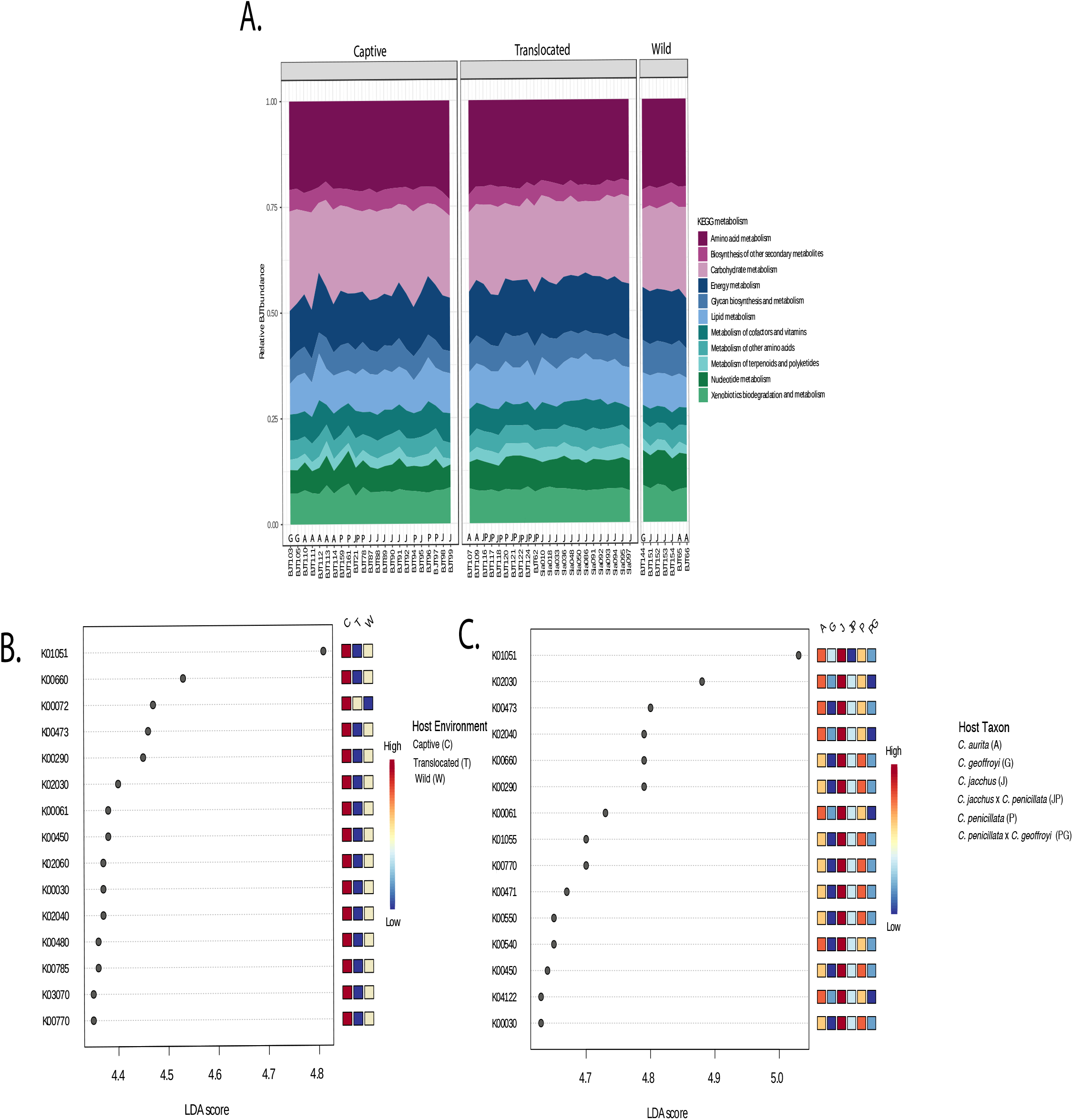
(A). Relative abundance of predicted KEGG pathways by host environment and taxon classification. (B). LefSe analysis of predicted KEGG pathway abundance by host environment. (C.) LefSe analysis of predicted KEGG pathway abundance by host taxon. Legend of host classifications are shown to the right of each plot.

Linkage analysis between *Callithrix* gut bacterial taxa and predicted gut microbiome function are show in Figures 6 and Supplementary Figures S3 and S4 for major bacterial and functional classes. Visual inspection of the three figures shows overall that different sets of bacterial taxa are responsible for carrying out different gut functional activities between captive and wild marmosets. Actinobacteria take on a number of functional roles in the *Callithrix* gut microbiome almost exclusively within wild hosts (Supplementary Figure S3). *Bifidobacterium* seems especially important among wild marmosets for carbohydrate and amino acid metabolism (Figure 6A). On the other hand, Proteobacteria seem to be heavily involved across variable major functions in the gut of captive and translocated marmosets (Supplementary Figure S3). Enterobacteriaceae seem to be carrying out a large number of functional roles, including across major categories of metabolic pathways in captive and translocated marmosets (Figure 6B). For other classes of bacteria found in the *Callithrix* gut, Bacteroidota, Firmicutes, and Campilobacterota seem to take on a broad number of functional roles in both translocated and captive marmosets (Supplementary Figure S4). The latter two also seems to also perform broad gut functional roles in a smaller subset of captive marmoset hosts (Supplementary Figure S4).

**Figure 6.**
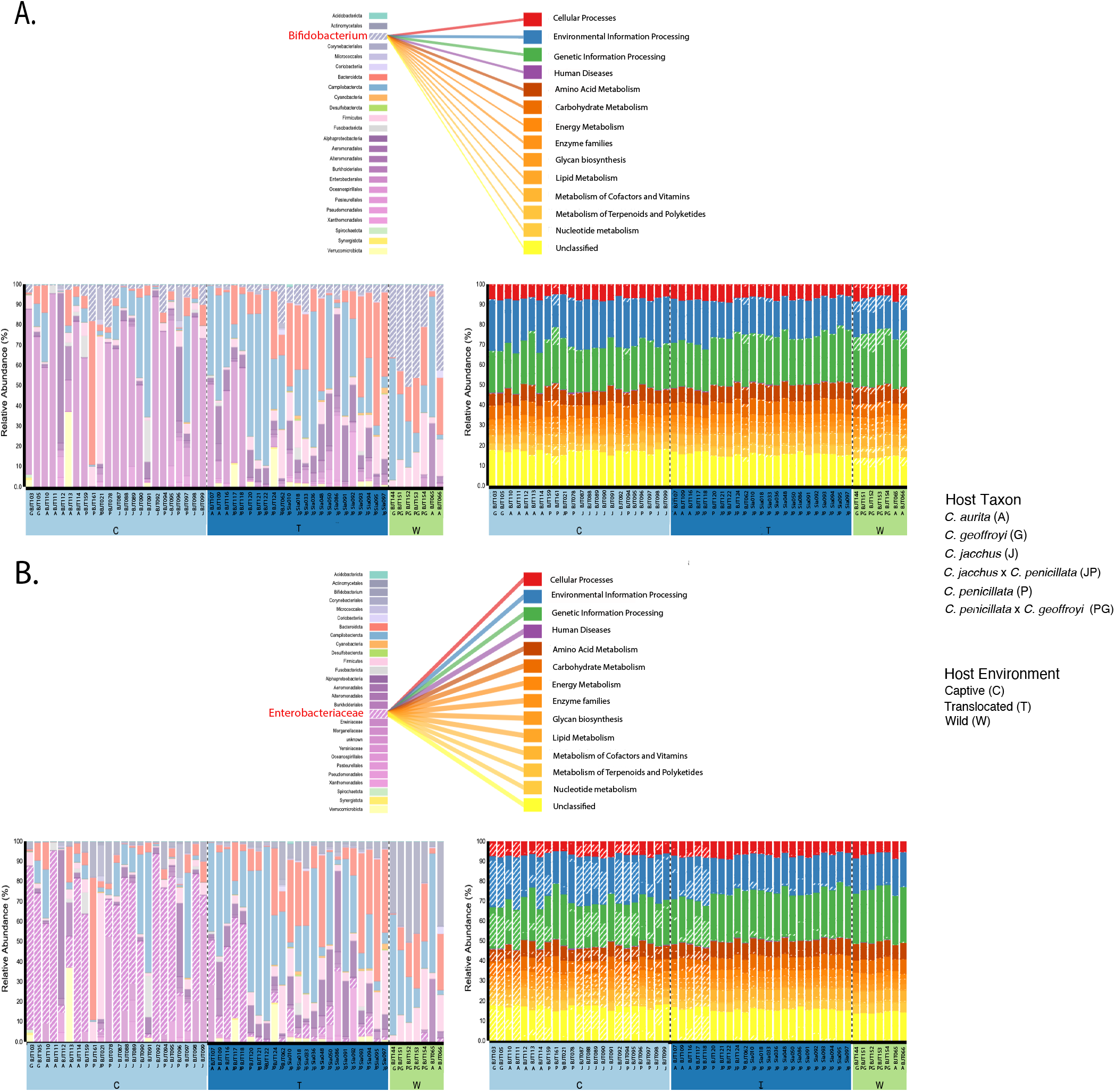
Visualization of BURRITO results showing linkage between *Callithrix* gut bacterial taxa composition and predicted functional profiles. In each plot, the lower left corner shows bacterial taxa relative abundance and the lower right shows predicted relative abundances of major functional categories of the *Callithrix* gut. The middle upper portion of each plot shows distribution of involvement of specific bacterial taxa in functional processes. Thickness of connecting lines between bacterial classes and functional classes indicates stronger involvement of a given bacterial taxon in a given functional process. The position of bacterial taxa and functional processes among respective relative abundance plots is represented by diagonal stripes. Host environment classifications in all plots are classified by C=Captive, T=Translocated, and W=Wild. (A) Distribution of *Bifidobacterium* role in predicted functional processes, with expansion of metabolic processes. (B) Distribution of Enterobacteriaceae role in predicted functional processes, with expansion of metabolic processes. Legend of host classifications are shown to the right of the plots.

## Discussion

In terms of *Callithrix* gut microbiome community structure, we found host taxon to significantly influence alpha diversity but not beta diversity. The significant pairwise differences in gut microbiome richness between *C. jacchus, C. penicillata*, and other marmosets may be related to relative differences for exudivory specialization between *Callithrix* taxa^31^. For example, *C. aurita* showed the lowest gut microbiome richness and is relatively one of the less specialized *Callithrix* taxa for exudate consumption^31^. In contrast, *C. jacchus* and *C. penicillata* are relatively the most specialized marmosets for gumnivory^31^, and possessed the highest levels of gut microbiome richness. A recent study of wild lemurs found that microbiomes, metagenomes, and metabolomes were species-specific and attuned to host dietary specializations and associated gastrointestinal morphology^68^. For *Callithrix*, a similar systematic study of taxa along a sliding scale of evolutionary specialization for exudivory is necessary to be undertaken with wild marmosets to better understand how host phylogeny influences gut microbiome diversity.

Host environment had nearly an overall significant effect on the *Callithrix* gut microbiome diversity. Significant differences were found between all host environmental classes for marmoset gut microbiome beta diversity, which is a result observed in various other animals (e.g., kiwis^15^, Tasmanian devil^17^, mice^16^, primates^5^, raptors^69^, rhinos^18^, woodrats^70^). A unique aspect of our study was the inclusion of hosts translocated from the wild into captivity, which were also significantly different from wild and captive hosts in terms of gut microbiome alpha diversity. In a similar vein, the gut microbiome of captive Tasmanian devils translocated into the wild exhibited temporal changes in gut microbiome diversity in response to the host’s changing environmental conditions^17^. Translocated hosts seem to importantly represent a dynamic transitional state between the relative extremes of wild and captive environments, which induce changes in host gut microbiome diversity. Overall, previous studies agree that dietary differences between host captive and wild environments are one of the main factors driving some of these gut microbiome changes.

In our sample, the gut microbiome of wild *Callithrix* was significantly enriched for the bacterial class of Actinobacteria, especially *Bifidobacterium*, and high abundance of *Bifidobacterium* in the gut microbiome may be a key biomarker for host gut microbiome eubiosis in marmosets^41^. This bacterial genus was also observable in the gut microbiome of captive and translocated marmosets we sampled, but to a much lesser degree. Across primates, *Callithrix* along with the closely related *Leontopithecus* are the two primate genera with the highest average abundance of *Bifidobacterium* (>30%) in the primate gut microbiome, followed by members of the Hominidae family (10%)^71^. While our sample size of wild marmosets was smaller relative to the number of captive and translocated/marmosets, our results are nonetheless the first to show that *Bifidobacterium* seem to be an integral part of the wild *Callithrix* gut microbiome. However, we were not able to determine the exact species of *Bifidobacterium* present in the gut of wild marmosets. Thus, an important next step in marmoset microbiome studies will be to expand study of wild marmosets and resolve wild *Callithrix* gut microbiome composition at the bacterial species level. Phylosymbiosis represents one promising approach to address this issue, as it combines genomic input data in the form of host phylogenetic markers or whole genomes and microbiome phylogenetic marker or meta-omics data^7^.

Several studies suggest that *Bifidobacterium* is a key component of the *Callithrix* gut microbiome to support carbohydrate metabolism^71–73^. In captive *C. jacchus*, species of *Bifidobacteria* in the gut microbiome were specific to host taxon and provided metabolic functions in line with *C. jacchus*’ relatively extreme adaptation to exudivory^73^. The *Bifidobacteria* group is especially efficient at metabolizing carbon sources like arabinogalactan and pectin^71^, which are components of carbohydrates of plant gums consumed by marmosets^74^. The genomes of three isolates of *Bifidobacterium callitrichos* from a captive *C. jacchus* fecal sample contained predicted genes associated with galactose and arabinose metabolism, which are also major constituents of tree gums eaten by *C. jacchus*^75^. In 3 US captive facilities, *C. jacchus* collectively shared four species of *Bifidobacterium*, which possessed genes encoding ATP-binding cassette proteins important for nutrient transport that may be specific to the marmoset gut^72^.

From this and previous studies, the gut microbiome composition of captive marmosets shows similarity to certain aspects of the gut microbiome composition of human gastrointestinal diseases associated with gut microbiome dysbiosis^4,27,41,76^. In our sample, the captive marmoset gut microbiome was overwhelming enriched for the Gammaproteobacteria bacterial class, and in particular from the family Enterobacteriaceae. In patients of Crohn’s Disease, the gut microbiome composition is enriched for bacterial taxa that include Enterobacteriaceae and depleted for Bifidobacteriaceae^27,77^. A similar pattern was observed in captive *C. jacchus* with gastrointestinal disease, which show various changes such as lowered *Bifidobacteria* abundance, rise in *Clostridium sensu stricto*, and the presence of Enterobacteriaceae in the cases of marmosets with inflammatory bowel disease^41^. Enterobacteriaceae is frequently associated with intestinal diseases and contains a number of pathogenic bacterial strains of Salmonella, Escherichia, and Shigella^78,79^. Perhaps the presence of Enterobacteriaceae in healthy captive marmosets makes them more susceptible for developing eventual gastrointestinal problems, as this is a shift away from the eubiosis of natural marmoset gut microbiome composition.

Translocated marmoset gut microbiome composition shows similarity to that of captive marmoset in being significantly enriched for the Proteobacteria phylum. However, translocated hosts possess a greater diversity of bacteria taxa within this phylum, as opposed to the higher gut Enterobacteriaceae abundance in captive hosts. One enriched Proteobacteria genus of note in the gut of translocated *Callithrix* was *Helicobacter*, of which certain species like *H. pylori* are known to cause gastric disease in humans^80^. Another Proteobacteria genus which was enriched in the translocated marmoset gut was *Campylobacter*. This bacterial genus is associated with diarrhea illness in humans^81^. Bacteroidetes and Clostridia were significantly abundant in the gut of translocated marmosets, a pattern also seen in the human gastrointestinal disease of ulcerative colitis^77^.

Despite the differences in gut bacterial composition and abundance among the marmosets in our sample, their gut microbiome seems to perform the same set of broad functions. Carbohydrate and amino acid metabolism are among the major functions carried out by the *Callithrix* gut microbiome in this study. However, closer inspection shows differential abundance of specific KEGG pathways between marmoset hosts from different environments. Further, there seems to be a stark difference in the distribution of functional roles among bacterial taxa found in the gut microbiome in captive, translocated, and wild marmosets. A relatively wider diversity of bacterial taxa take on functional roles in the gut microbiome of translocated and wild marmosets. *Bifidobacterium* seems to take a prominent role in amino acid and carbohydrate metabolism in wild marmosets, a pattern not replicated in non-wild marmosets. Instead, in captive and translocated marmosets, Proteobacteria seem to dominate functional roles of the gut microbiome. In captive marmosets, Enterobacteriaceae seem to dominate all aspects of gut microbiome function.

Given that gastrointestinal distress is highly prevalent in captive dietary specialist NHPs^21–24^, several authors suggest that host dietary specialization and its direct connection with the gut microbiome is an important factor affecting health outcomes of captive hosts. Certain bacteria are selected for in the gut according to the energetic substrates available from the host diet^82^, thus designing captive diets need to be planned carefully^83^. The chemical composition of sugars in the marmoset diet have be most explored in *Callithrix jacchus*, and include beta-linked polyssachrides composed on galactose, arabinose, and rhamnose^75^. Additionally, pectin is another carbohydrate found in the bark of *Anadenanthera peregrina*, which is consumed by various taxa of marmosets^74^. Wild marmosets also generally exploit other nutritional sources such as fruit, fungi, and small prey^31,84^. In captivity, marmosets diets do not reflect what marmosets would normally eat in nature. When gum is supplied to marmosets in captivity, the most commonly used source is arabica gum^42^. However, most captive institutions do not supplement marmoset diets with gum, and instead they generally combine different proportions of commercial chow, fruits, vegetables, protein, and sweets^42^. Captive Brazilian facilities where we sampled marmosets for this study also follow similar husbandry practices for marmoset nutrition (Table 4) as that described by Goodroe et al^42^.

**Table 4.** Diet collectively fed to marmoset hosts in sampled captive facilities.

|  |  |
| --- | --- |
| Fruits | Papaya, Orange, Banana, Apple, Pear, Avacado, Kiwi, Melon, Mango |
| Carbs | Sweet Potato, Potato, Beets |
| Vegetables | Cucumber, Eggplant, Pumpkin, Chuchu, Cauliflower, Carrots |
| Proteins | Cooked Chicken, Cooked Egg |

One concern for a lack of tree gums in the diet of captive specialist exudivores is the development of health issues as well as a negative impact on breeding and survivability^85,86^. In humans suffering from gastrointestinal diseases, increasing plant-based foods and dietary fiber, resulted in increasing microbiome diversity, remission of gastrointestinal symptoms, and decreasing risk of gastrointestinal distress^4,87^. Such diets may increase gut abundance of bacteria such as *Bifibacterium* that produce short chain fatty acids like butyrate, which may guard against proliferation of pathogenic bacteria in the gut and decrease chronic inflammation^4,87^. Lack of access to a natural-wild diet for marmosets and other exudivory specialists may also promote loss of native gut microbes like *Bifidobacterium* and enrichment of potentially pathogenic bacterial strains of Enterobacteriaceae. It has also been demonstrated in mice and wood-rats that feeding more natural diets to individuals in captivity helped maintain host gut microbiome composition profiles closer to wild, free-ranging hosts^88,89^. One noteworthy study of folivorous captive sifakas carried out systematic experimental dietary manipulations while integrating metagnomics and metabolomics data to determine how foliage quality affected gut microbiome composition and production of colonic short chain fatty acids. We stress that similar studies need to be undertaken for marmosets and specialized exudivores^23^. There is especially a need to determine if provisioning of gum in the diet of captive exudivores will lead to improved host welfare by maintaining gut microbiomes closer to that of wild populations.

Our major study findings are consistent with previous studies in showing that gut microbiome composition is sensitive to host environmental factors, and that *Bifidobacterium* may be an important biomarker for marmoset gut microbiome health. We also show that carbohydrate metabolism is a key function of the *Callithrix* gut microbiome. Given our limited sampling of wild marmosets, further studies with expanded sampling of wild individuals representing all *Callithrix* species are still needed. For exudivores in general, more studies are needed to understand better the health and reproductive consequences of omitting as well as increasing gum intake by specialized exudivores in captivity. Overall, such information will expand baseline gut microbiome data available for wild and non-wild exudivores to allow for the development of new tools to improve exudiviore management, welfare, and conservation.

## Acknowledgements

We thank Vanner Boere, Ita de Oliveira, Rodrigo S. Carvalho, the Guarulhos Zoo, the CEMAFAUNA, the CPRJ staff, SERCAS staff, and AMLD staff for assistance with wild and captive populations. We are very grateful to Cecil M. Lewis and Tanvi Honap at LMAMR for donation of PCR primers. We thank Dietmar Zinner, Corinna Ross, Tauras Vilgalys, and four anonymous reviewers for comments on this work.

## Author contributions statement

JM and ADG formulated the idea for the study. JM collected samples, obtained funding, conducted wet and dry laboratory work, and wrote the original manuscript. RAC provided study guidance, and logistical support. JAD, SK, LTN, ATO provided study guidance, and logistical support. CSI gave access and provided logistical support to collect samples from animals kept at Guarulhos Zoo. SBM provided logistical support and veterinary assistance to collect samples from animals kept at the Rio de Janeiro Primatology Center. PAN gave access and provided logistical support to collect samples from animals kept at CEMAFAUNA. AAQ provided logistical support and veterinary assistance to collect samples from animals kept at CEMAFAUNA. LCMP gave access and provided logistical support to collect samples from animals kept at CEMAFAUNA. AP gave access and provided logistical support to collect samples from animals kept at CPRJ. CRRM was a major contributors in writing the manuscript and provided access to collect samples from SERCAS marmosets. DLS performed DNA extractions, methodological optimization, and carried out PCR. ACS was a major contributor in writing the manuscript, provided study guidance, and logistical support. CR gave significant guidance and input in the development of this study. All authors read and approved the final manuscript.

## Additional information

The dataset supporting the conclusions of this article is available in the NCBI SRA repository under Bioproject PRJNA574641. The authors declare that they have no competing interests.

