## Supplemental Figure Legends for "The Gut Microbiome of Exudivorous Marmosets in the Wild and Captivity"

Supplementary Figure S1. Sample rarefaction curve after rarefying for the sample with the lowest sequencing depth.

Supplementary Figure S2. (a) LefSe analysis of bacterial class abundance categorized by host taxon. (b) LefSe analysis of bacterial genus abundance categorized by host taxon.

Supplementary Figure S3. Visualization of BURRITO results showing linkage between *Callithrix* gut bacterial taxa composition and predicted functional profiles. In each plot, the upper left corner shows bacterial phylogeny, the lower left corner shows bacterial taxa relative abundance, the upper right shows a network of functional processes, and the lower right shows predicted relative abundances of major functional categories of the *Callithrix* gut. The middle upper portion of each plot shows distribution of involvement of specific bacterial taxa in functional processes. Thickness of connecting lines between bacterial classes and functional classes indicates stronger involvement of a given bacterial taxon in a given functional process. The position of bacterial taxa and functional processes among respective relative abundance plots is represented by diagonal stripes. Host environment classifications in all plots are classified by C=Captive, T=Translocated, and W=Wild. (a). Distribution of Actinobacteriota role in predicted functional processes. (b) Distribution of Proteobacteria role in predicted functional processes.

Supplementary Figure S4. Visualization of BURRITO results showing linkage between *Callithrix* gut bacterial taxa composition and predicted functional profiles. In each plot, the upper left corner shows bacterial phylogeny, the lower left corner shows bacterial taxa relative abundance, the upper right shows a network of functional processes, and the lower right shows predicted relative abundances of major functional categories of the *Callithrix* gut. The middle upper portion of each plot shows distribution of involvement of specific bacterial taxa in functional processes. Thickness of connecting lines between bacterial classes and functional classes indicates stronger involvement of a given bacterial taxon in a given functional process. The position of bacterial taxa and functional processes among respective relative abundance plots is represented by diagonal stripes. Host environment classifications in all plots are classified by C=Captive, T=Translocated, and W=Wild. (a) Distribution of Acidobacteriota role in predicted functional processes. (b) Distribution of Bacteroidota role in predicted functional processes. (c) Distribution of Campilobacterota role in predicted functional processes. (d) Distribution of Cyanobacteria role in predicted functional processes. (e) Distribution of Desulfobacterota role in predicted functional processes. (f) Distribution of Firmicutes role in predicted functional processes. (g) Distribution of Fusobacteriota role in predicted functional processes. (h) Distribution of Spirochaetota role in predicted functional processes. (i) Distribution of Synergistota role in predicted functional processes. (j) Distribution of Verrucomicrobiota role in predicted functional processes.
