## Supplementary figures and images for "The Gut Microbiome of Exudivorous Marmosets in the Wild and Captivity"

### Supplementary Figure S1

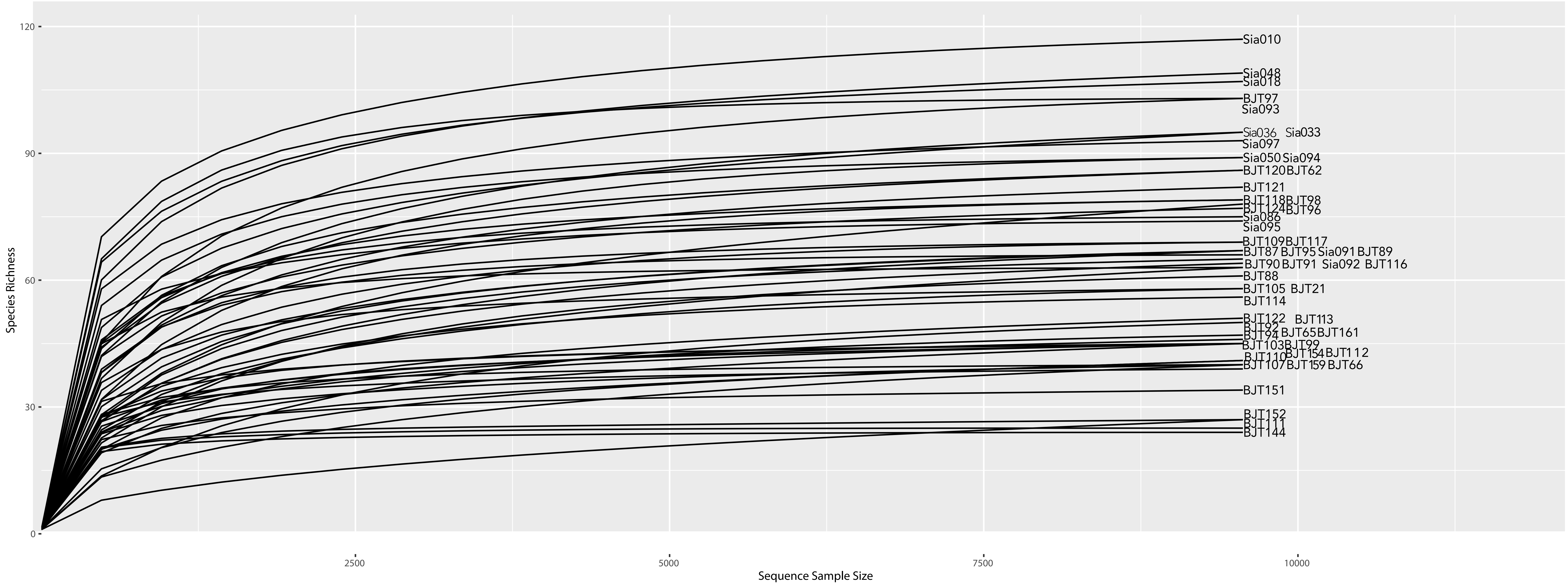

### Supplementary Figure S2

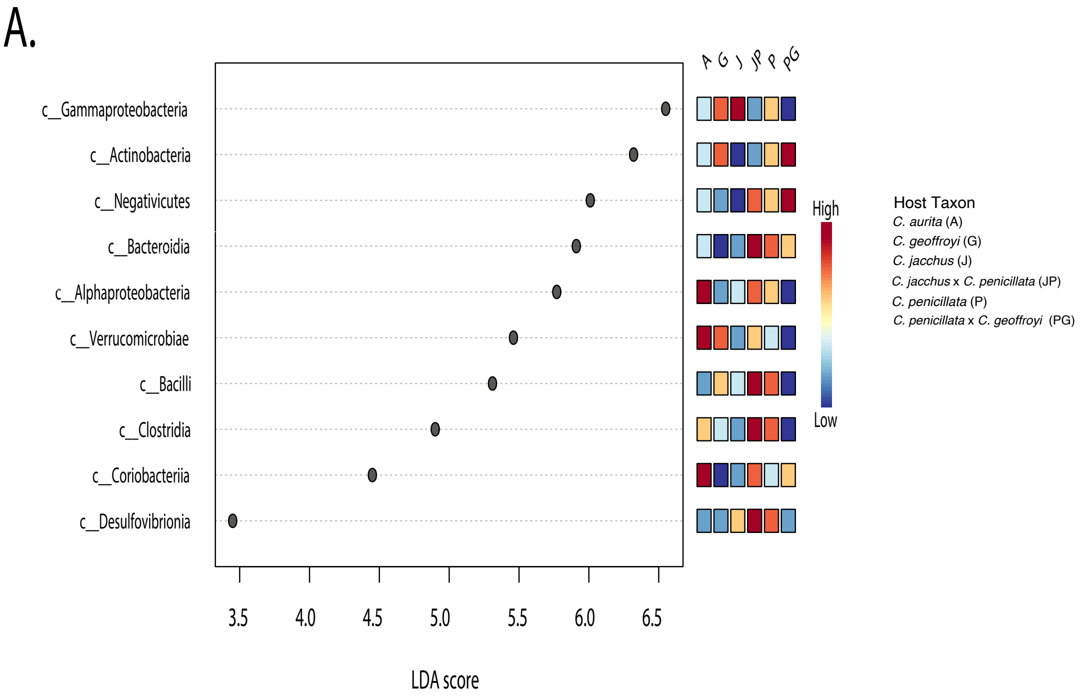

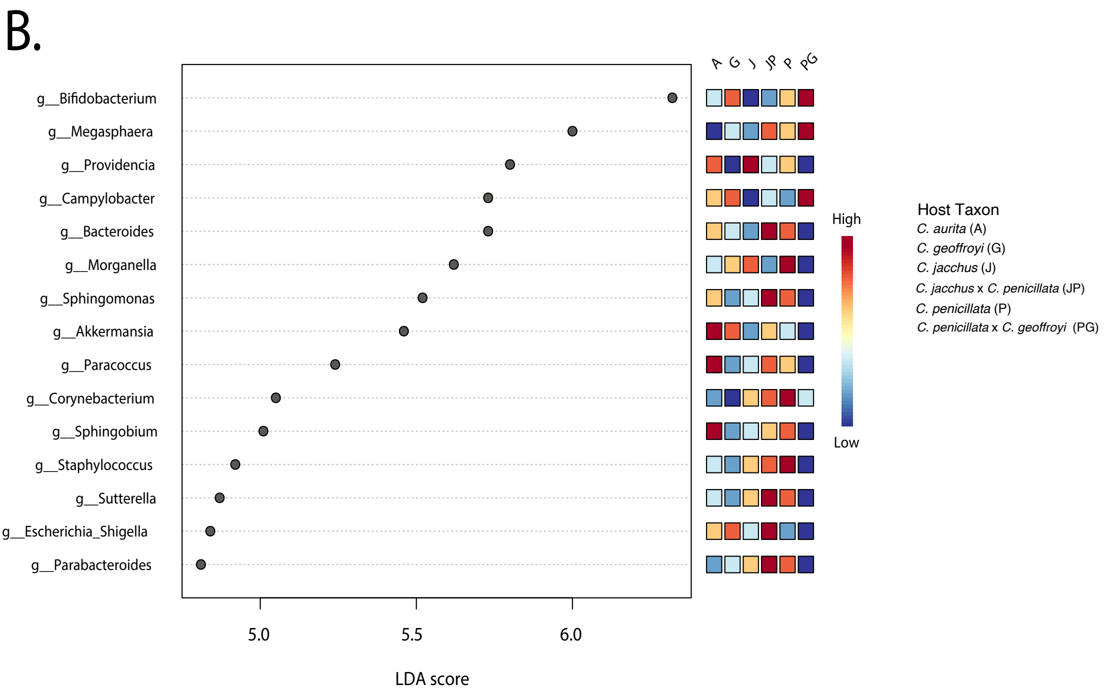

### Supplementary Figure S3

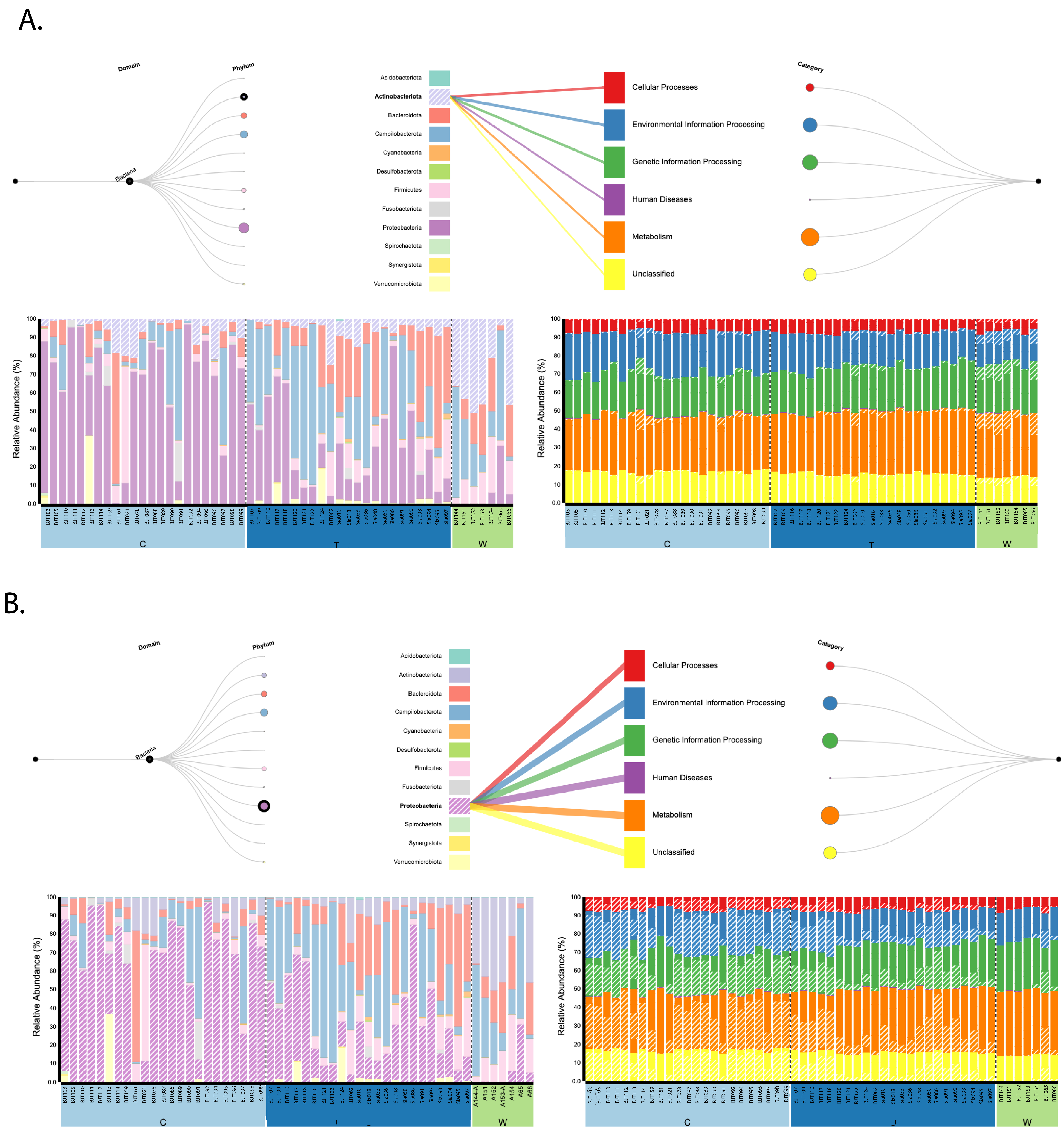

### Supplementary Figure S3

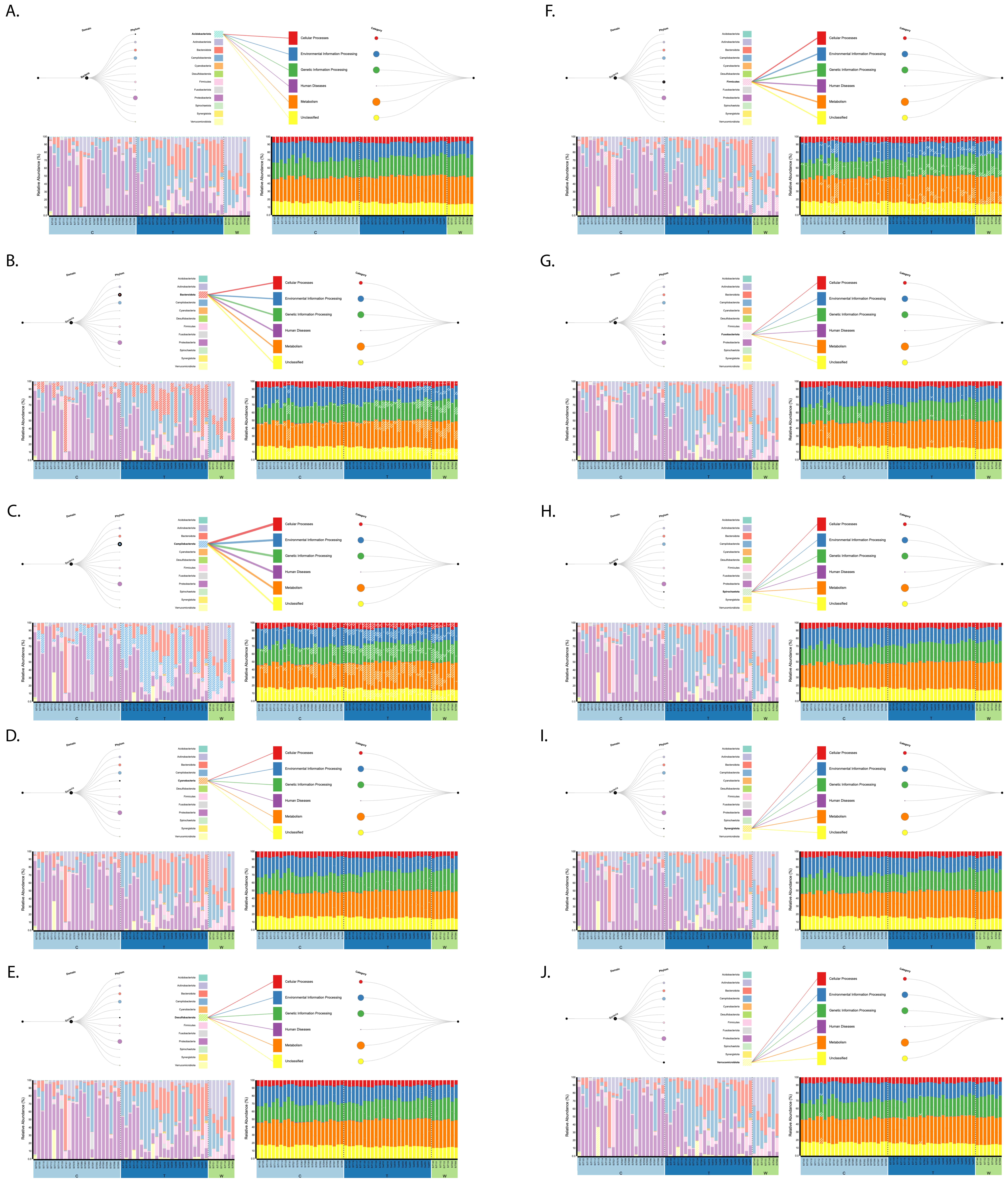
